# Mutant SF3B1 promotes PDAC malignancy through TGF-β resistance

**DOI:** 10.1101/2022.06.16.496393

**Authors:** Patrik T. Simmler, Tamara Mengis, Kjong-Van Lehmann, André Kahles, Tinu Thomas, Gunnar Rätsch, Markus Stoffel, Gerald Schwank

**Affiliations:** Institute of Molecular Health Sciences, ETH Zurich, 8093 Zurich, Switzerland; Institute of Pharmacology and Toxicology, University of Zurich, 8057 Zurich, Switzerland; Department of Computer Science, ETH Zurich, 8092 Zurich, Switzerland; Swiss Institute of Bioinformatics, Zurich, Switzerland; Department of Biology, ETH Zurich, 8093 Zurich, Switzerland; University Hospital Zurich, 8091 Zurich, Switzerland

**Keywords:** SF3B1, K700E mutation, splicing, pancreatic cancer, TGF-β, apoptosis, MAP3K7

## Abstract

The splicing factor SF3B1 is recurrently mutated in various tumors, including pancreatic ductal adenocarcinoma (PDAC). The impact of the hotspot mutation SF3B1^K700E^ on the PDAC pathogenesis, however, remains elusive. Here, we demonstrate that Sf3b1^K700E^ alone is insufficient to induce malignant transformation of the murine pancreas, but increases aggressiveness of PDAC if it co-occurs together with mutated KRAS and p53. We further demonstrate that SF3B1^K700E^ reduces epithelial–mesenchymal transition (EMT) and confers resistance to TGF-β1-induced cell death, and provide evidence that this phenotype is in part mediated through aberrant splicing of *Map3k7*. Taken together, our work suggests that SF3B1^K700E^ acts as an oncogenic driver in PDAC through enhancing resistance to the tumor suppressive effects of TGF-β.

## INTRODUCTION

Genes involved in RNA splicing are frequently mutated in various cancer types (Yoshida et al., 2011). The splicing factor subunit 3b 1 (SF3B1) is amongst the most commonly mutated components of the splicing machinery, with high incidence in myelodysplastic syndromes (MDS) (Je et al., 2013) and chronic lymphocytic leukemia (CLL) (Miao et al., 2019). However, also in various solid tumors SF3B1 is recurrently mutated, including uveal melanoma (UVM) (Furney et al., 2013), breast cancer (BRCA) (Fu et al., 2017; Maguire et al., 2015; Sun et al., 2020), prolactinomas (Li et al., 2020) hepatocellular carcinoma (HCC) (Zhao et al., 2021) and pancreatic adenocarcinoma (PDAC) (Bailey et al., 2016; Yang et al., 2021). As part of the U2 small nuclear ribonucleoprotein (U2 snRNP) SF3B1 exerts an essential function in RNA splicing by recognizing the branchpoint sequence (BPS) of nascent RNA transcripts (Wahl et al., 2009; Zhang et al., 2020). This process is crucial for the definition of the 3’ splice site (3’ ss) of the upstream exon-intron boundary, a prerequisite for the accurate removal of introns (Wahl et al., 2009). It is well understood that hotspot mutations in SF3B1 at HEAT repeats 5-9 allow the recognition of an alternative BPS, resulting in the inclusion of a short intronic region into the mature messenger RNA (mRNA) (Alsafadi et al., 2016; Canbezdi et al., 2021; Darman et al., 2015; DeBoever et al., 2015; Kesarwani et al., 2017). These alternatively spliced transcripts are prone to degradation through nonsense mediated RNA decay (NMD) (Darman et al., 2015). Several recent studies have evaluated the mechanistic contribution of genes mis-spliced by oncogenic SF3B1 to tumor progression. So far, missplicing of *PPP2R5A* was found to increase malignancy through stabilizing c-Myc (Liu et al., 2020a; Yang et al., 2021), and aberrant *MAP3K7* splicing was reported to promote NF-κB–driven tumorigenesis (Liu et al., 2021).

Despite compelling evidence on the oncogenic role of mutated SF3B1 in hematologic malignancies, its contribution to the formation and/or progression of solid tumors is less well understood. Since splicing deregulation has been reported as a hallmark of PDAC, with SF3B1 being recurrently mutated (Bailey et al., 2016), we aimed at elucidating the impact of the frequently occurring SF3B1^K700E^ mutation to the pathogenesis of this tumor type. We demonstrate that *Sf3b1^K700E^* increases malignancy in a mouse model for PDAC by decreasing the sensitivity of tumor cells to TGF-β-induced cell-death. We further provide evidence that TGF-β-resistance is mediated through missplicing of *Map3k7*. Together, our work suggests that SF3B1^K700E^ exerts its oncogenic role in PDAC by dampening the tumor-suppressive effect of TGF-β.

## RESULTS

### SF3B1^K700E^ is a tumor driver in PDAC

RNA processing has been previously identified as a hallmark of pancreatic cancer in the PACA-AU cohort (Bailey et al., 2016). Validating these findings, we found that genes encoding for the splice factors RBM10, SF3B1 and U2AF1 are also frequently mutated in the PACA-CA cohort (Suppl. Fig. 1A). In accordance with its described function as a tumor suppressor (Hernández et al., 2016), 56% of the mutations found in RBM10 lead to a truncated protein. Conversely, in SF3B1 and U2AF1 the majority of mutations were missense mutations that occurred at hotspot sites, indicating a neomorphic function of the mutated proteins. Like in other cancer types, also in PDAC the most frequently found mutation in SF3B1 led to a lysine (K) to glutamic acid (E) amino acid change at position 700 (SF3B1^K700E^) (Suppl. Fig. 1B). Therefore, we experimentally tested if the SF3B1^K700E^ mutation contributes to PDAC malignancy by generating a mouse model where the *Sf3b1^K700E^*mutation is specifically activated in the pancreas using Ptf1a-Cre (Fig. 1A, Suppl. Fig. 1C). First, we tested if *Sf3b1^K700E/+^* alone could already induce PDAC formation. However, heterozygous activation of the *Sf3b1^K700E^*allele did not have any effect on survival of mice or weight of the pancreas after 300 days (Fig. 1B, C). In addition, no difference in tissue architecture and cell proliferation was observed (Suppl. Fig. 1D, E). Next, we assessed if the K700E mutation could enhance aggressiveness of PDAC, and crossed the *Sf3b1^K700E^* allele into the *Ptf1a-Cre; Kras^G12D/+^; Trp53^fl/fl^* (KPC) mouse model (Fig. 1D), which is known to induce PDAC within 2 to 3 months (Bardeesy et al., 2006; Hingorani et al., 2005; Marino et al., 2000). Importantly, KPC-Sf3b1^K700E/+^ animals displayed a significantly shorter survival and an increased tumor size compared to KPC mice, although no obvious differences in tissue architecture were found (Fig. 1E-G, Suppl. Fig. 1F-H). Likewise, we also observed an effect on tumor malignancy when we introduced the *Sf3b1^K700E/+^* mutation into *Kras^G12D/+^* mice, which is a model for pre-cancerous pancreatic neoplasms with sporadic PDAC formation after a prolonged latency period (Hingorani et al., 2005). Introducing the *Sf3b1^K700E/+^* mutation in *Kras^G12D/+^* mice resulted in increased pancreas weight, a larger area of neoplastic pancreas tissue (Fig. 1I - K), and in 64% of mice compared to 10% of *Kras^G12D/+^* control mice pancreatic lesions developed into PDAC at an age of 43 weeks (Fig 1L).

**Figure 1.**
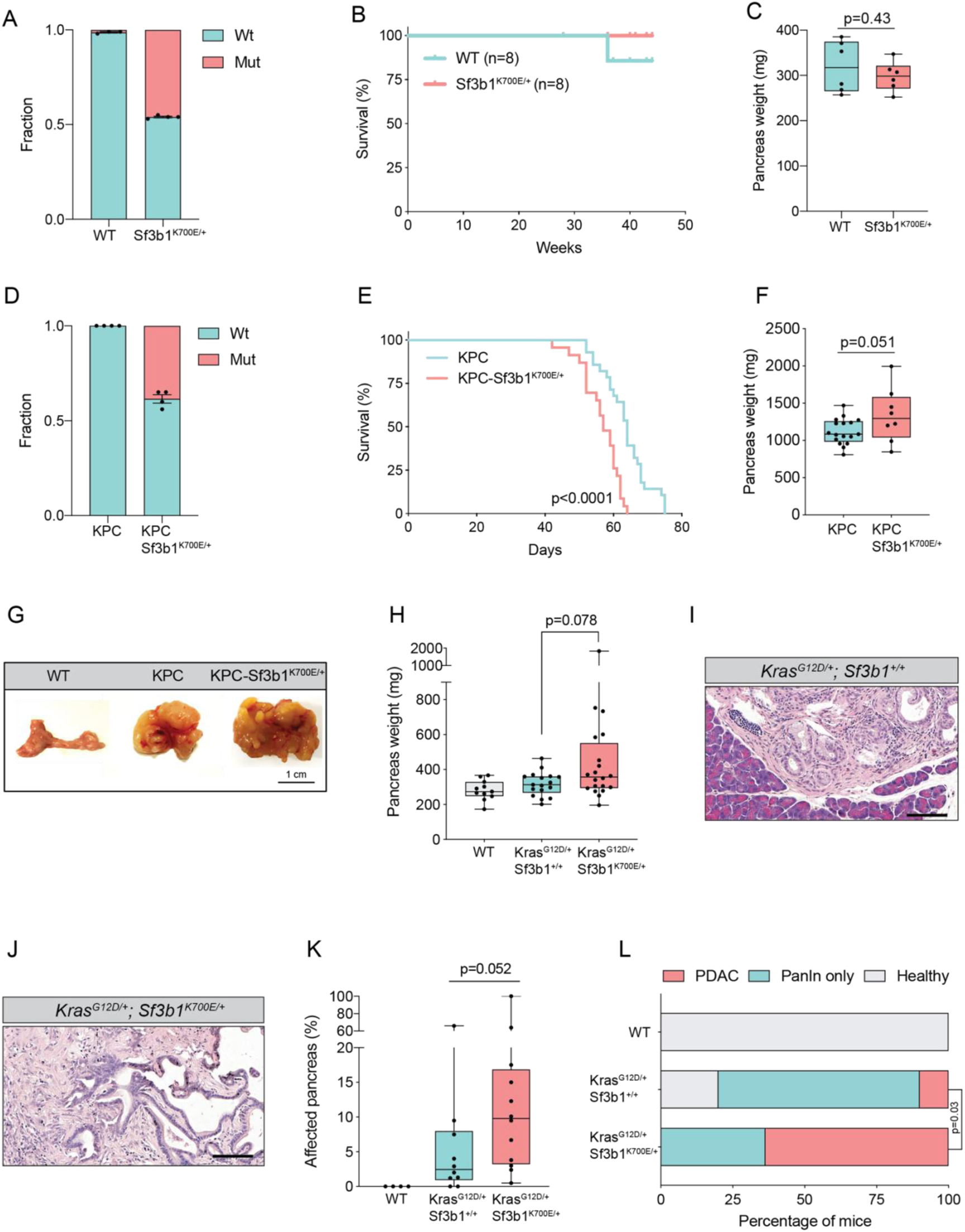
Sf3b1^K700E^ increases aggressiveness of murine PDAC. **(A)** Fraction of the K700E mutation (T>C) of cDNA isolated from pancreata of *Ptf1a-Cre* (WT) (n=3) or *Ptf1a-Cre; Sf3b1^K700E/+^* mice (n=4), assessed by Sanger-sequencing. **(B)** Survival of WT and *Sf3b1^K700E/+^* mice followed over 300 days (n=8). **(C)** Pancreatic weight of WT and *Sf3b1^K700E/+^*mice at 300 days of age (3 males and 3 females for each genotype). Two-tailed unpaired t-test was used to compute the indicated p-value. **(D)** Fraction of K700E mutation (T>C) of RNA isolated from of sorted KPC (n=3) and KPC-Sf3b1^K700E/+^ cells (n=4), assessed by RNA-seq. **(E)** Survival of KPC and KPC-Sf3b1^K700E/+^ mice. P-value was determined by Log-rank (Mantel-Cox) testing. **(F)** Pancreatic weight of KPC and KPC-Sf3b1^K700E/+^ mice at 9 weeks of age. Two-tailed unpaired t-test was used to compute the indicated p-value (* p<0.05). **(G)** Representative photographs of WT, KPC and KPC-Sf3b1^K700E/+^ pancreata. **(H)** Pancreatic weight of WT, *Kras^G12D/+^* and *Kras^G12D/+^; Sf3b1^K700E/+^* mice at 43 weeks of age. Welch’s unequal variances t-test was used to compute the indicated p-value. **(I-J)** Representative micrograph images of *Kras^G12D/+^* (I) and *Kras^G12D/+^; Sf3b1^K700E/+^* (J) pancreata at 43 weeks of age stained with H&E, scale bar is 100 μm. **(K)** Affected area (including PanINs and PDAC) of WT, *Kras^G12D/+^* and *Kras^G12D/+^; Sf3b1^K700E/+^* pancreata at 43 weeks of age. Mann-Whitney test was used to compute the indicated p-value. **(L)** Percentage of mice at 43 weeks of the indicated genotypes showing PanINs (blue) or PanINs and PDAC formation (red). P-value indicates significance of the difference in PDAC formation, computed by Chi-square test.

### SF3B1^K700E^ reduces expression of EMT genes in pancreatic tumors

In order to elucidate the functional impact of SF3B1^K700E^ on the transcriptome, we isolated cancer cells of mouse tumors by fluorescence-activated cell sorting (FACS) of Epithelial cell adhesion molecule (EpCAM) positive cells and performed RNA-sequencing (RNA-seq) (Table 1). High purity of isolated tumor cells was confirmed by the absence of sequencing reads for *Trp53* exons 2–10, which are excised via Cre-recombination specifically in tumor cells (Suppl. Fig. 2A), and by the presence of the *Sf3b1^K700E^* mutation in 38% of the transcripts (Fig. 1D). Principal component analysis separated the sequenced replicates (3 KPC and 4 KPC-Sf3b1^K700E/+^ tumors) according to the genotype, indicating a major impact of the K700E mutation on the transcriptome (Suppl. Fig. 2B).

We next performed gene set enrichment analysis (GSEA), which revealed IFN-α-response as the most significantly enriched pathway in KPC-Sf3b1^K700E/+^ tumor cells (Suppl. Fig. 2C). This result is in line with a previous study, which found that aberrant splicing caused by SF3B1 inhibition or oncogenic SF3B1 mutations induces an IFN-α-response through retinoic acid-inducible gene I (RIG-I) mediated recognition of cytosolic aberrant RNA-species (Chang et al., 2021). Interestingly, the most significantly depleted gene set in KPC-Sf3b1^K700E/+^ cells was epithelial-mesenchymal transition (EMT) (Fig. 2A, B, Suppl. Fig. 2C). We first confirmed downregulation of the most significantly depleted gene of the EMT gene set, the glycoprotein Tenascin-C (*Tnc)*, by qPCR on additional KPC-Sf3b1^K700E/+^ tumor samples (Fig. 2C) and by histology in KPC-Sf3b1^K700E/+^PDAC sections (Fig. 2D, E). Next, we assessed if the reduction of EMT genes was induced cell-autonomously by the *Sf3b1^K700E/+^* mutation, or if it was an indirect consequence of the altered micro-environment in *Sf3b1^K700E^* KPC tumors. We therefore compared the expression of the 15 most significantly depleted EMT genes (Fig. 2B) *in vitro* in *Sf3b1^K700E^* vs. *Sf3b1* WT KPC pancreatic organoids. Importantly, 71% of the analysed genes were significantly reduced in KPC-Sf3b1^K700E/+^ vs. KPC organoids, with none of the genes showing a trend towards elevated expression (Fig. 2F). Finally, we analysed whether differences in EMT gene expression could be a consequence of differences in the tumor stage between *Sf3b1^K700E^* versus *Sf3b1* WT KPC tumors. We therefore established non-cancerous pancreatic organoids from LSL-*Kras^G12D/+^; Trp53^fl/fl^; Sf3b1^flK700E/+^* and LSL-*Kras^G12D/+^; Trp53^fl/f^; Sf3b1*^+/*+*^ mice, and induced recombination *in vitro* through lentiviral Cre transduction (Suppl. Fig. 2D). However, also in this experimental setup, 67% of the analysed EMT genes were significantly downregulated in *Sf3b1^K700E^* vs. *Sf3b1* WT organoids, with only one of the analysed genes showing a minor trend for elevated expression (Suppl. Fig. 2E). Together, these data indicate that *Sf3b1^K700E^*mediates downregulation of EMT genes in a cell autonomous manner independently of the PDAC microenvironment or stage.

**Figure 2.**
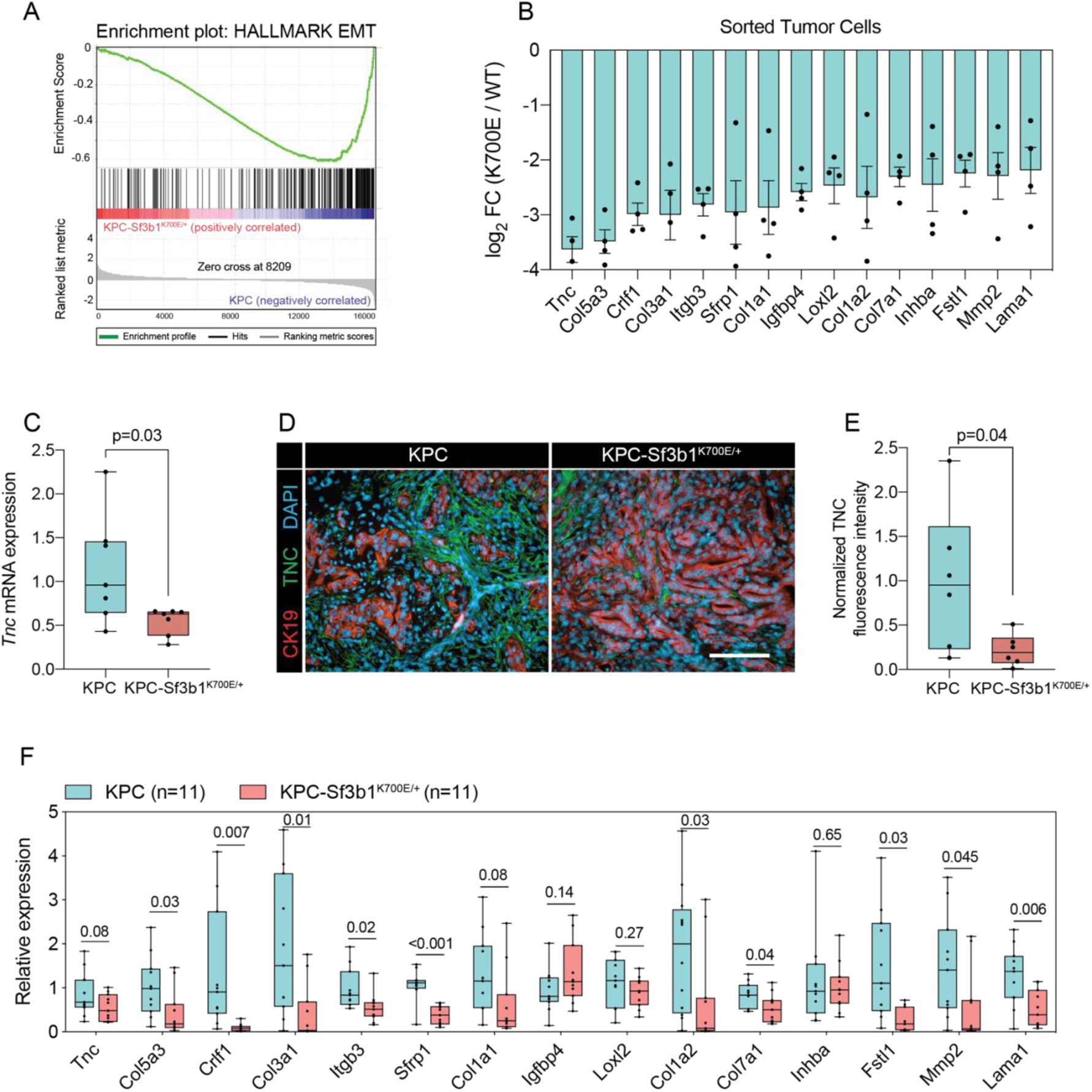
*Sf3b1^K700E^* induces downregulation of EMT. **(A)** Gene-set enrichment analysis (GSEA) enrichment plot of epithelial-mesenchymal transition (EMT), representing the most deregulated pathway of the GSEA-Hallmark pathways when comparing KPC (n=3) and KPC-Sf3b1^K700E/+^ (n=4) sorted tumor cells. **(B)** Top 15 of downregulated genes of the GSEA-EMT gene list in sorted KPC-Sf3b1^K700E/+^ cells (FDR < 0.05, logCPM > 1). **(C)** *Tnc* expression in KPC (n=7) and KPC-Sf3b1^K700E/+^ (n=7) tumors, assessed by RT-qPCR. Two-tailed unpaired t-test was used to compute the indicated p-value. **(D)** Representative Immunofluorescence staining of CK19 (red) and TNC (green) in murine PDAC samples, counterstained with DAPI (blue). Scale bar is 50 μm. **(E)** Quantification of TNC staining in KPC (n=6) and KPC-Sf3b1^K700E/+^ (n=6) tumors. The averaged area of TNC staining in 3 randomly chosen fields per tumor specimen was compared by a two-tailed unpaired t-test. **(F)** The expression of the EMT genes displayed in (B) was assessed by RT-qPCR in tumor-derived cancer cell lines (KPC, n=11 and KPC-Sf3b1^K700E/+^ n=11) after one passage of ex-vivo culture. For analysis, Ct-values of the indicated genes were normalized to *Actb* and a two-tailed unpaired t-test was used to compute the indicated p-values.

### SF3B1^K700E^ confers resistance to TGF-β1-induced cell death

The two major EMT-promoting cytokines are TNF-α and TGF-β (Bulle and Lim, 2020). We therefore determined if the EMT genes that are most significantly downregulated by *Sf3b1^K700E^* in KPC tumours are induced by TNF-α or TGF-β. Importantly, we found that 80% of the analysed genes were strongly induced by TGF-β in tumour derived KPC cells (Fig 3A), whereas TNF-α significantly upregulated only 20% of these genes (Suppl. Fig. 2F). Furthermore, we observed that *in vitro* induced *Sf3b1^K700E^*KPC organoids show a 6-fold reduced invasion through matrigel when stimulated with TGF-β, linking the repression of TGF-β responsive EMT genes with a functional migratory impairment (Figure 3B, C).

**Figure 3.**
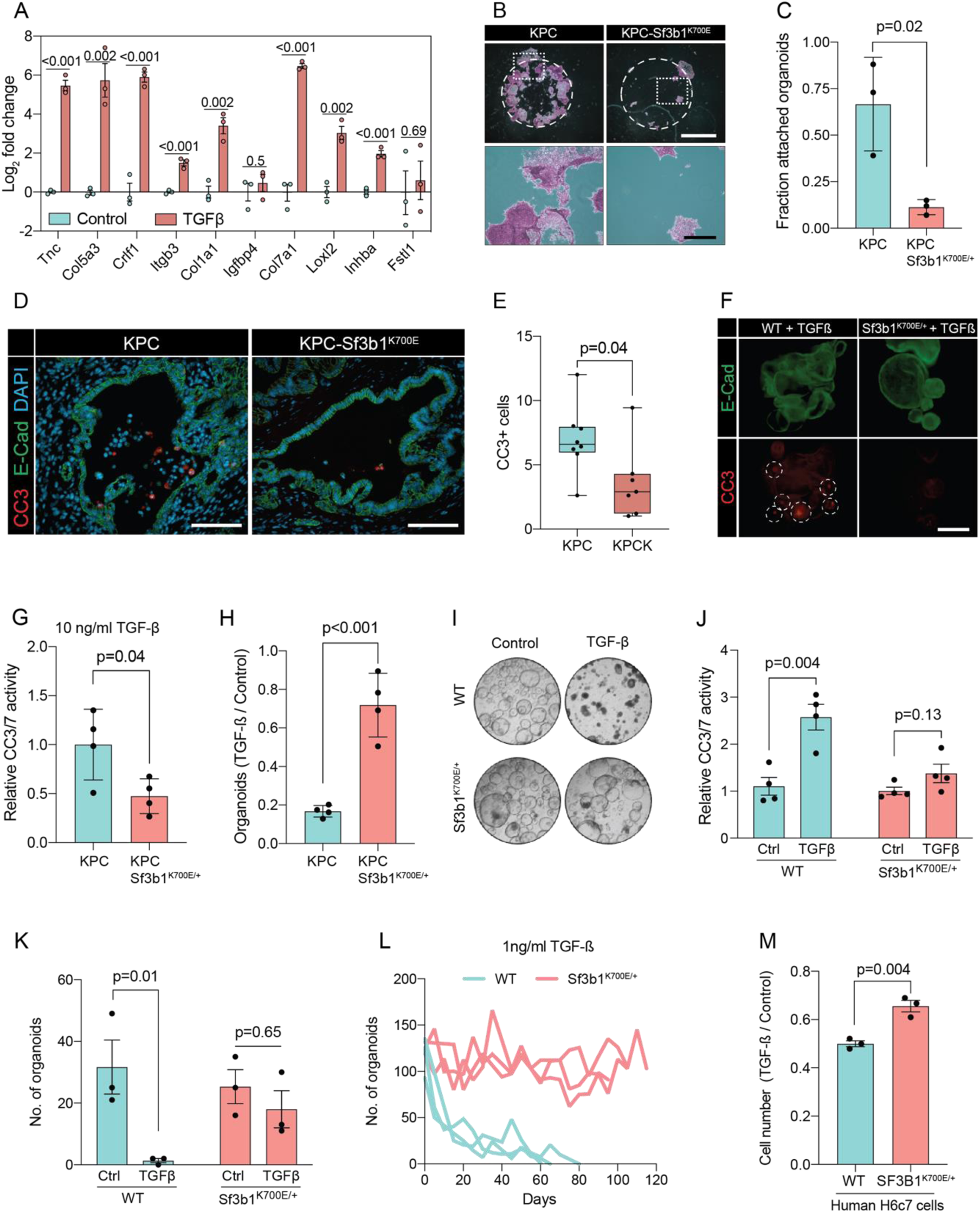
*Sf3b1^K700E^* reduces TGF-β-induced apoptosis. **(A)** RT-qPCR analysis of EMT genes displayed in Fig. 2B in 3 different KPC cell lines treated with 10 ng/ml TGF-β1 for 24 hours. The experiment was performed independently 3 times for every cell line. *Col3a1, Sfrp1, Col1a2, Mmp2* and *Lama1* were not detected and therefore excluded from analysis (see methods for details). **(B)** Representative micrographs of KPC (n=3) and KPC-Sf3b1^K700E/+^ (n=3) cancer organoid lines treated with TGF-β1 (10 ng/ml) for 48 hours. Matrigel was detached prior to staining with crystal violet. Scale bar is 1 mm (panel above) or 100 μm (panel below). **(C)** Quantification of (B). The fraction of attached organoids was calculated by dividing the number of attached organoids by the number of total organoids. The experiment was performed independently 3 times for every cell line, the average of all replicates is shown. Two-tailed unpaired t-test was used to compute the indicated p-value. **(D)** Representative microscopy images of E-Cadherin (green) and CC3 (red) in murine PDAC samples. Scale bar is 50 μm. **(E)** Quantification of CC3 positive cells in KPC (n=8) and KPC-Sf3b1^K700E/+^ (n=7) tumor samples. The average number of CC3 positive cells of 5 microscopic fields is plotted, two-tailed unpaired t-test was used to compute the indicated p-value. **(F)** Immunofluorescence staining of E-Cadherin (green) and Cleaved Caspase 3 (CC3, red) in WT and Sf3b1^K700E/+^ organoids exposed to TGF-β1 (10 ng/ml) for 12 hours. CC3 positive cells are highlighted by white dashed lines. Scale bar is 100 μm. **(G)** Quantification of Cleaved Caspase 3 and 7 (CC3/7) activity measured by Caspase-Glo assay of KPC (n=3) and KPC-Sf3b1^K700E/+^ (n=4) in-vitro activated cancer cell lines treated with TGF-β1 (10 ng/ml) for 24 hours. The experiment was repeated independently twice for every cell line, the average of the replicates is shown. Two-tailed unpaired t-test was used to compute the indicated p-value. **(H)** Quantification of viable organoids of the indicated genotype exposed to 10 ng/ml TGF-β1 for 48 hours, normalized to organoid numbers of untreated control samples. Each data point shows a different organoid line. For each organoid line, the experiment was independently performed three times, the average of replicates is plotted. Two-tailed unpaired t-test was used to compute the indicated p-value. **(I)** Representative microscopy images of WT and *Sf3b1^K700E/+^* organoids exposed to 10 ng/ml TGF-β1 for 48 hours. **(J)** Quantification of CC3/7 in WT and *Sf3b1^K700E/+^* organoids exposed to 10 ng/ml TGF-β1 for 48 hours. The experiment was repeated independently four times. Two-tailed unpaired t-test was used to compute the indicated p-values. **(K)** Quantification of viable organoids of the indicated genotype exposed to 10 ng/ml TGF-β1 for 48 hours, normalized to organoid numbers of untreated control samples. Each data point shows a different organoid line. For each organoid line, the experiment was independently performed three times, the average of replicates is plotted. Two-tailed unpaired t-test was used to compute the indicated p-values. **(L)** Organoid count of organoids cultured in medium containing 1 ng/ml TGF-β1 for up to 120 days. One organoid line per genotype was used, the experiment was repeated three times independently. **(M)** Viability of the human pancreatic duct cell line H6c7 overexpressing wildtype or mutated SF3B1 after 72 hours of exposure to 10 ng/ml TGF-β1 assessed by crystal violet staining. The optical density of TGF-β1 treated cells was normalized to untreated control cells. The experiment was independently performed three times, two-tailed unpaired t-test was used to compute the indicated p-value.

In pancreatic lesions TGF-β induces EMT, followed by apoptosis of the affected cells in a process termed lethal EMT (David et al., 2016). This prompted us to speculate that SF3B1^K700E^ could drive PDAC progression by reducing sensitivity of epithelial cells to TGF-β-mediated lethal EMT. Performing immunofluorescence staining for cleaved caspase 3 in KPC tumors, we first confirmed that the majority of apoptotic cells reside in the lumen of PanINs (Suppl. Fig. 2G) and that these cells are negative for the epithelial markers E-cadherin and high Fibronectin-1 (Fig. 3D, Suppl. Fig. 3A) (Hruban et al., 2006). We then analysed *Sf3b1^K700E^* tumors by immunofluorescence staining. In line with our hypothesis, we observed a reduction in luminally extruded cells and a reduction in cleaved caspase 3-positive cells (Fig. 3D, E, Suppl. Fig. 3B). To further analyse the impact of *Sf3b1^K700E^* on the tumor suppressive effect of TGF-β, we exposed *Sf3b1* WT and *Sf3b1^K700E^* KPC tumor organoids to TGF-β1. We again observed reduced caspase 3 and 7 activity in *Sf3b1* mutant organoids, and a greatly increased survival rate (72% vs. 17% surviving organoids, Fig. 3F-H). Since the tumor-suppressing effect of TGF-β is most prominent on pre-cancerous epithelial cells (Massagué, 2008), we additionally established organoid lines with- and without *Sf3b1^K700E^* from non-cancerous mouse pancreata. While *Sf3b1^K700E^*reduced proliferation without supplementation of TGF-β1 (Suppl. Fig. 3C), treatment with TGF-β1 led to significantly lower caspase 3 and 7 activity in *Sf3b1^K700E/+^* organoids, and to significantly enhanced survival (77% vs. 3%) (Fig. 3I-K). Likewise, also in *Kras^G12D/+^* organoids SF3B1^K700E^ led to increased survival in the presence of TGF-β1 (Suppl. Fig. 3D). Finally, exposure to low levels (1 ng/ml and 2 ng/ml) of TGF-β1 also allowed long-term expansion of *Sf3b1^K700E/+^* organoids (analysed for over 120 days), while the number of *Sf3b1* WT organoids rapidly declined within the first 15 days (Fig. 3L, Suppl. Fig. 3E, F).

To assess if reduced sensitivity to TGF-β is also observed in human pancreas cells containing the SF3B1^K700E^ mutation, we stably overexpressed either wildtype or mutated SF3B1 in the pancreatic duct cell line H6c7. This cell line is derived from healthy pancreatic tissue, which unlike all tested PDAC-derived cell lines (BxPC-3, Mia PaCa-2, PANC-1 and PSN-1) is still partially responsive to the suppressive effect of TGF-β signalling (Fig. 3M, Suppl. Fig. 3G-K). In line with our results from murine PDAC, overexpression of mutant SF3B1^K700E^ resulted in an increased viability upon TGF-β1 exposure compared to overexpressing wildtype SF3B1 (Fig. 3M), while no effect of SF3B1^K700E^ on proliferation was observed in absence of TGF-β1 treatment in human pancreatic duct cells and PDAC cell lines (Suppl. Fig. 3L).

### SF3B1^K700E^ reduces TGF-β sensitivity through *Map3k7* missplicing

To identify how SF3B1^K700E^ could mediate TGF-β resistance, we next assessed the impact of the mutation on RNA-splicing. By analysing RNA-seq data from sorted KPC vs. KPC-Sf3b1^K700E/+^ tumor cells we predominantly identified alternative 3’ splice events (Fig. 4A, Table 2), with cryptic 3’ ss showing an upstream adenosine enrichment and a less pronounced polypyrimidine tract most often located 8-14 bases upstream of the canonical 3’ ss (Fig. 4B, C). These findings are in accordance with previous splice-analyses performed in various murine SF3B1 mutant tissues (Liu et al., 2021, 2020a; Mupo et al., 2017; Obeng et al., 2016; Yin et al., 2019). Next, we sought to determine which of the identified K700E-dependent alternative splice events are conserved between mice and humans. Due to limited publicly available RNA-seq datasets in human PDAC, we analysed a pan-cancer dataset containing samples of 32 different cancer types. In agreement with previous studies, we found that also in human cancers the SF3B1^K700E^ hotspot mutation leads to a predominant use of cryptic 3’ ss (Suppl. Fig. 4A, B, Table 2) (Alsafadi et al., 2016; DeBoever et al., 2015; Kesarwani et al., 2017; Tang et al., 2020; Wang et al., 2016). Importantly, we further identified 11 genes that contained an alternative 3’ splice-event linked to SF3B1^K700E^ in human tumors and KPC mice (Fig. 4D). Of those genes, *MAP3K7* (formerly known as TGF-β activated kinase 1 / *TAK1*) in particular raised our attention. It is a well-described effector of cytokine-signalling that mediates non-canonical TGF-β signalling (Kim and Choi, 2012), and it was recently shown to induce EMT and apoptosis in TGF-β stimulated human mammary cells (Tripathi et al., 2019). Using targeted RNA-seq, we found that one third of *Map3k7* transcripts were misspliced in pancreata of *Sf3b1^K700E/+^* and KPC-Sf3b1^K700E/+^ mice (Fig. 4E, F, Suppl. Fig. 4C). Confirming inter-species conservation of this alternative splice-event, we also found *MAP3K7* missplicing in H6c7 cells and human PDAC cell lines overexpressing SF3B1^K700E^ (Fig. 4F, Suppl. Fig. 4D). Notably, we also assessed SF3B1^K700E^ dependent alternative splicing of *Ppp2r5a*, which was reported to impair apoptosis via post-translational modification of BCL2 in leukaemia (Liu et al., 2020a). However, we did not observe significant alternative 3’ss usage or mRNA expression of *Ppp2r5a* in *Sf3b1^K700E^* mutant KPC tumors (Suppl. Fig. 4E, F), indicating tissue-specificity of this splice-event.

**Figure 4.**
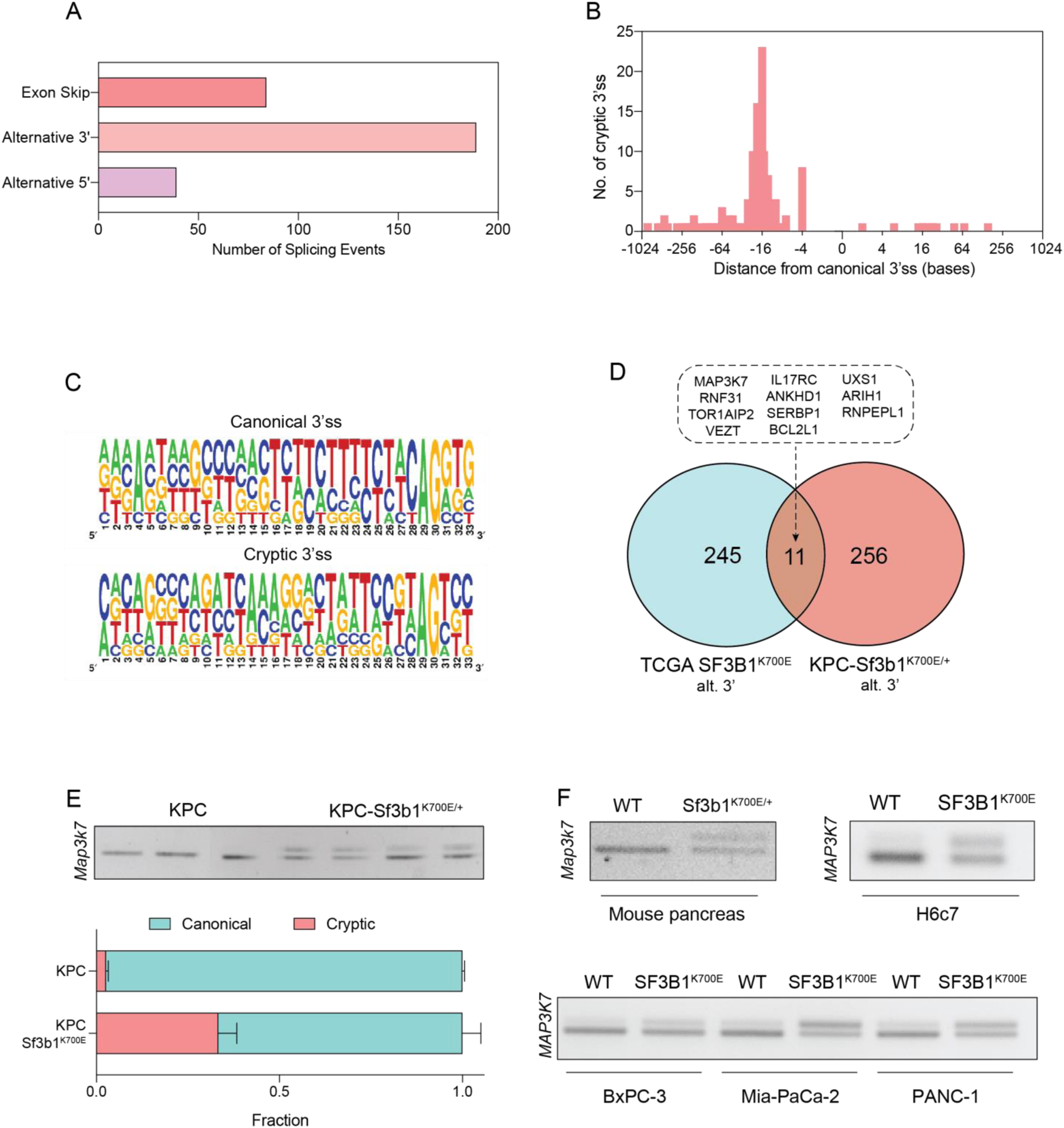
SF3B1-K700E predominantly induces aberrant 3’ splice site selection. **(A)** Summary of alternative splice events detected in KPC-Sf3b1^K700E/+^ sorted tumor cells (PSI > 0.1, p < 0.01). **(B)** Histogram displaying the distance of cryptic 3’ss from the adjacent canonical 3’ss in sorted KPC-Sf3b1^K700E/+^ tumor cells on a logarithmic scale. **(C)** Consensus 3’ ss motif in proximity of the canonical (top) and the cryptic (bottom) 3’ ss for 7 alternative 3’ splicing events identified in sorted KPC-Sf3b1^K700E/+^ tumor cells. **(D)** Venn-diagram depicting alternative 3’ splice events in the pan-cancer dataset (PSI>0.05 and p<1^-10^) and sorted KPC-Sf3b1^K700E/+^ tumor cells (PSI>0.1, p<0.01). **(E)** Representative gel image (top) and NGS-results (bottom) of *Map3k7* cDNA isolated from sorted KPC and KPC-Sf3b1^K700E/+^ tumor cells (n=3) (A). The amplicon includes the 3’ splice site of exon 4 and 5, the upper band of the gel image represents the non-canonical transcript variant. **(F)** Representative gel image of RT-PCR amplicon of *Map3k7* cDNA isolated from WT and *Sf3b1^K700E/+^* pancreata and from the indicated human cell lines.

Since *Sf3b1^K700E^* dependent missplicing in *MAP3K7* was shown to result in reduced RNA and protein levels of MAP3K7 in leukaemia (North et al., 2022), we hypothesized that the reduced responsiveness to TGF-β signalling in SF3B1^K700E^ mutant pancreas cells is caused by lower MAP3K7 levels. First, we confirmed a reduction in Map3k7 levels in vitro and in vivo in *Sf3b1^K700E^* mutant pancreatic cells by RT-qPCR and western blotting (Fig. 5A-D). Next, we tested whether this reduction could explain the observed resistance to TGF-β in *Sf3b1*-mutant PDAC, and assessed the expression of EMT genes in TGF-β treated KPC cells with a stable knock-down of MAP3K7 (Fig. 5E, Suppl. Fig. 4G). A decrease of *Map3k7* mRNA levels to 35% (SD ± 10%) led to a reduced expression in 7 out of 10 EMT genes (Suppl. Fig. 4H). Knocking down *Map3k7* in pancreatic organoids, moreover, led to increased viability upon TGF-β1-treatment (Fig. 5F, Suppl. Fig. 4I), and chemical inhibition of p38, one of the major effectors of MAP3K7, partially protected organoids against TGF-β1 induced cell death (Fig. 5G). Further supporting our hypothesis that *Sf3b1^K700E^* mediates resistance to TGF-β1 via MAP3K7, overexpression of the full-length isoform of MAP3K7 in TGF-β1-treated *Sf3b1^K700E^* mutant organoids significantly decreased their viability (Fig. 5H, I).

**Figure 5.**
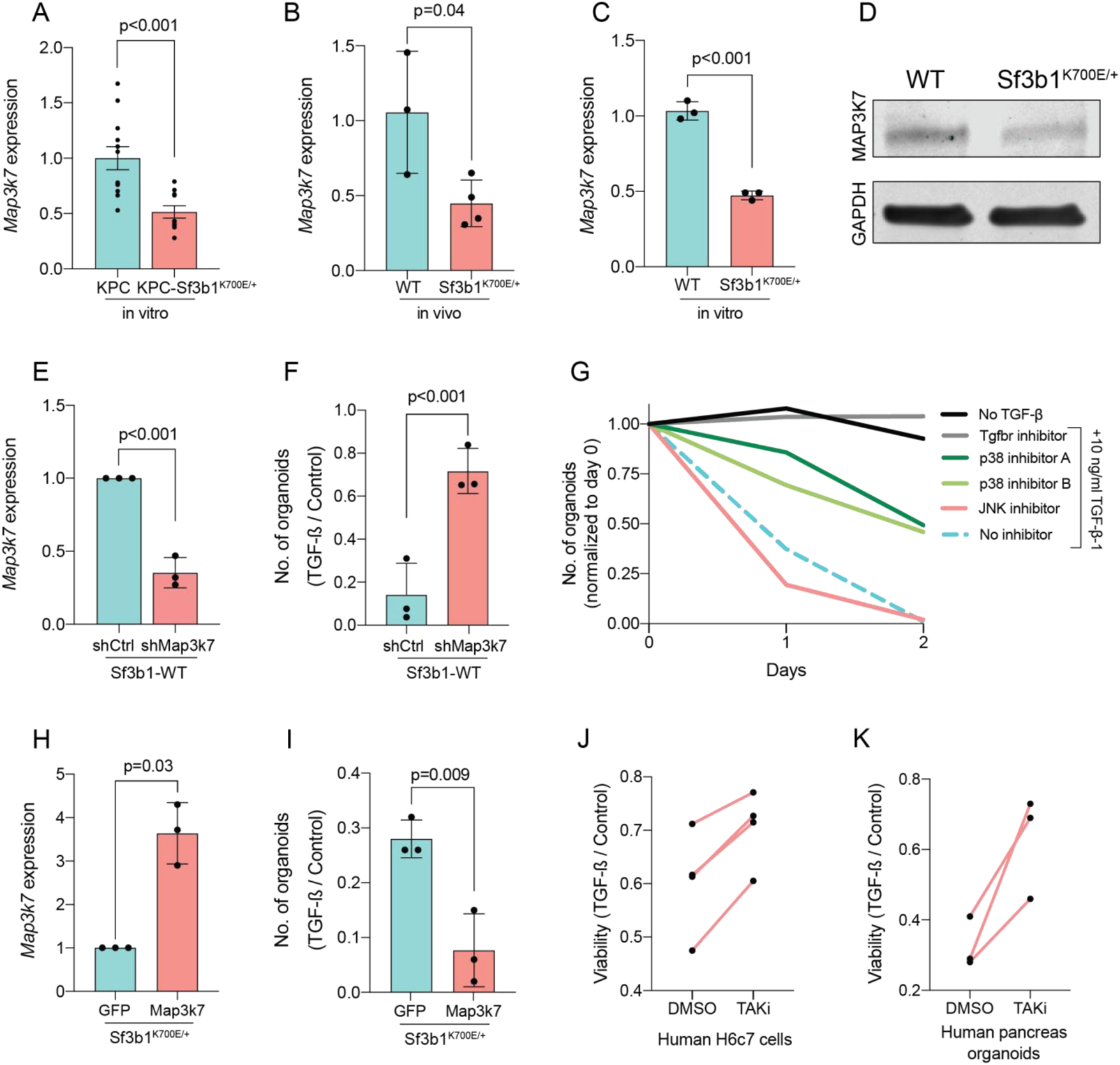
Reduction in *Map3k7* lowers sensitivity to TGF-β1. **(A-C)** RT-qPCR data showing *Map3k7* expression in KPC (n=13) and KPC-Sf3b1^K700E/+^ (n=12) cancer-derived organoid lines (A), as well as WT (n=3) and Sf3b1^K700E/+^ (n=4) pancreata (B) and organoid lines (C). For analysis, Ct-values of *Map3k7* were normalized to *Actb* and a two-tailed unpaired t-test was used to compute the indicated p-values. Data show mean and standard error of the mean in A and B. **(D)** Representative Western blot gel-image of MAP3K7 and GAPDH in WT and Sf3b1^K700E/+^ organoids. **(E)** RT-qPCR analysis of *Map3k7* in cells transduced with a shMap3k7 compared to a control shRNA, a two-tailed unpaired t-test was used to compute the indicated p-values. **(F)** Quantification of viable WT and *Sf3b1^K700E/+^* murine pancreatic duct organoids or WT transduced with control shRNA (shCtrl) or shRNA targeting *Map3k7* (shMap3k7). The organoids were exposed to 10 ng/ml TGF-β1 for 24 hours prior to analysis. The experiment was independently performed three times. Two-tailed unpaired t-test was used to compute the indicated p-value. **(G)** Viability of wildtype (WT) organoids cultured in medium containing 10 ng/ml TGF-β1, supplemented with chemical inhibitors targeting the indicated effectors of TGF-β1-signalling. Two independent experiments are summarized. Further details are provided in the methods section. **(H)** RT-qPCR analysis of *Map3k7* in cells transduced by lentivirus with an overexpression construct of *Map3k7*, compared to overexpression of GFP, a two-tailed unpaired t-test was used to compute the indicated p-values. **(I)** Quantification of viable murine pancreatic duct organoids with stable overexpression of *Map3k7*, exposed to 10 ng/ml TGF-β1 for 96 hours. Data represents one organoid line per condition, the experiment was independently performed three times. Two-tailed unpaired t-test was used to compute the indicated p-value. **(J-K)** Viability of human pancreatic duct H6c7 cells (J) or human pancreatic duct organoids (K) exposed to 10 ng/ml TGF-β1 with addition of the MAP3K7 inhibitor Takinib (TAKi, 5 μM) or DMSO. The viability was assessed after 48 hours of TGF-β1 treatment and normalized to cells grown in absence of TGF-β1. The experiment was independently performed four (J) or three (K) times.

To finally assess conservation of this mechanism between mice and humans, we treated human pancreatic H6c7 cells as well as human pancreatic organoids exposed to TGF-β1 with Takinib, a chemical inhibitor for MAP3K7. In line with our results in murine cells, also in the human pancreatic cells inhibition of MAP3K7 led to a significant increase in survival (Fig. 5J, K). Taken together, our results suggest that *Sf3b1^K700E^* mediates resistance of pancreatic epithelial cells to TGF-β1 via MAP3K7, providing a potential mechanism for its role of in PDAC progression.

## DISCUSSION

The frequent occurrence of SF3B1 hotspot mutations in various tumor types implies a contribution to tumorigenesis. While the molecular function of oncogenic SF3B1 on RNA-splicing is well described, how deregulation of misspliced genes contribute to malignancy in different cancer entities is less understood. Previous studies have shown that in chronic lymphocytic leukaemia (CLL) SF3B1^K700E^ leads to missplicing of *PPP2R5A,* which in turn stabilizes c-Myc and thereby promotes aggressiveness of tumor cells (Liu et al., 2020a; Yang et al., 2021). Furthermore, in breast cancer SF3B1^K700E^ causes a tumor-promoting effect through missplicing in *MAP3K7* and downstream activation of NF-κB-signalling (Liu et al., 2021). In our study we analysed the oncogenic function of SF3B1^K700E^ in the context of PDAC. Using a mouse model, we provide the first experimental evidence that the mutation indeed promotes PDAC progression. We show that SF3B1^K700E^ reduces TGF-β induced EMT and cell-death, thereby providing a potential explanation for the oncogenicity of SF3B1^K700E^ in PDAC. In line with our hypothesis, pancreatic epithelial cells have previously been found to undergo lethal EMT when stimulated with TGF-β (David et al., 2016), and acquiring TGF-β resistance is considered to be essential in early stages of PDAC tumorigenesis (Hezel et al., 2012).

A PDAC-promoting effect of reduced TGF-β signalling was previously demonstrated in a conditional mouse model for *Smad4*, a core-component of TGF-β-signalling. In line with our observations, *Smad4* deletion was not sufficient to induce malignant transformation, but was found to be oncogenic only in combination with concomitant activation of Kras^G12D^ (Bardeesy et al., 2006b). SF3B1^K700E^, nevertheless, only partially suppressed non-canonical TGF-β-signalling, and the inactivation of canonical TGF-β-signalling via SMAD4 inactivation or both branches via TGF-β receptor inactivation is likely to confer additional tumor promoting effects (Hezel et al., 2012). Therefore, we speculate that the SF3B1^K700E^ mutation does not eliminate the selective pressure in PDAC for additional inactivating mutations in TGF-β-signalling, and in case additional mutations impede both branches of TGF-β signalling, SF3B1^K700E^ might lose its beneficial role for the tumor. Like in myeloid cancers, targeting SF3B1^K700E^ via small molecules might therefore only benefit a subset of patients carrying this mutation (Steensma et al., 2021).

Reducing *Map3k7* levels by shRNA did not impair TGF-β-mediated EMT and cell death to the same extend as *Sf3b1^K700E^*. It is therefore likely that the aberrant splicing of other genes is also contributing to the observed resistance to TGF-β. Nevertheless, our hypothesis that TGF-β resistance in SF3B1^K700E^ mutant PDAC is at least partly mediated via MAP3K7 is also supported by previous studies. These already demonstrated that SF3B1 mutations induce 3’ missplicing of *MAP3K7* in various tumor entities (Bondu et al., 2019; Li et al., 2021; Lieu et al., 2022; Liu et al., 2020b; Wang et al., 2016; Zhang et al., 2019), that aberrant splicing by SF3B1^K700E^ reduces MAP3K7 protein levels (North et al., 2022) and that MAP3K7 mediates TGF-β-induced EMT and apoptosis in mammary epithelial cells (David et al., 2016).

While the most common oncogenic drivers in PDAC are well described, oncogenes occurring at lower frequency are understudied (Hudson et al., 2010). This study provides a first demonstration that oncogenic SF3B1^K700E^ promotes tumor progression *in vivo* in a mouse model for PDAC, and sheds mechanistic insights on the oncogenicity of SF3B1^K700E^ by reducing the responsiveness to tumor-suppressive TGF-β.

## AUTHOR CONTRIBUTIONS

P.S. and G.S. conceived the study. G.S., M.S. and G.R. supervised the study. P.S. designed, executed and analyzed most of the in vitro and in vivo experiments. T.M. performed in vitro experiments. TT. and AK. developed and ran the RNA-seq pipeline, K.L. performed the in-silico statistical analysis of RNA-splicing in human and murine samples. P.S. and G.S. wrote the manuscript with input from all authors.

## ACKNOWLEDGEMENTS

We are grateful to E. A. Obeng (Dana-Farber Cancer Institute, Boston) and B. L. Ebert (Brigham and Women’s Hospital, Harvard Medical School, Boston) for providing the Sf3b1^K700E/+^ mice used in our study. We are also greatful to Prof. Wilhelm Krek (Department of Biology, Institute of Molecular Health Sciences, ETH Zurich, Switzerland). He initiated this project, but sadly passed away in August 2018. This work was financed by grants from the Swiss National Science Foundation. TT., AK., KL are supported by ETH core funding to Gunnar Rätsch.

## METHODS

### Animal models

*Sf3b1^K700E/+^* mice were a gift from E. A. Obeng (Dana-Farber Cancer Institute, Boston, USA) and B. L. Ebert (Brigham and Women’s Hospital, Harvard Medical School, Boston, USA). *LSL-Kras^G12D/+^*, *LSL-Trp53^R172H/+^*and *Ptf1a-Cre* mice were purchased from the Jackson Laboratory (Bar Harbor, Maine, USA). All *Sf3b1^K700E/+^* and KPC-*Sf3b1^K700E/+^* mice were bred in a C57BL/6J background, *Sf3b1^K700E/+^; Kras^G12D/+^* mice were a C57BL/6J-BALB/c strain. Female and male mice were used for all experiments. Animals displaying dwarfism were excluded from analysis. The minimum of animals needed for the study was estimated by Fisher-Yates analysis. Due to the observed variance of the mouse model, more mice than initially estimated were used for the study. Mice were held in a specific-pathogen-free (SPF) animal facility at the ETH Phenomics Center EPIC (ETH Zurich, Switzerland). All animal experiments were conducted in accordance with the Swiss Federal Veterinary Office (BVET) guidelines (license no. ZH055/17).

### Cell lines

The human cell lines AsPC-1 (CRL-1682), BxPC-3 (CRL-1687), MIA-PaCa-2 (CRM-CRL275 1420), PANC-1 (CRL-1469), PSN-1 (CRL-3211) and HEK293T (CRL-1573) cells were purchased from ATCC. H6c7 cells (ECA001-FP) were purchased from Kerafast. MIA-PaCa277 2, PANC-1 and HEK293T cells were maintained in DMEM with 4.5 g/l D-Glucose and GlutaMAX (Gibco), supplemented with 10% fetal calf serum (FCS, Sigma-Aldrich) and 1% Penicillin-streptomycin (P/S, Invitrogen). AsPC-1, BxPC-3 and PSN-1 were cultured in RPMI1640 (Thermo Fisher), supplemented with 10% FCS and 1% P/S. H6c7 cells were cultured in Keratinocyte serum-free medium (Thermo Fisher), supplemented with recombinant EGF and bovine pituitary extract according to the manufacturer’s instructions (Thermo Fisher), as well as 100 μg/ml Primocin (Invivogen). Cell lines were regularly checked for mycoplasma infections by Mycoplasma PCR-detection test (Thermo-Fisher).

### Murine organoids

Murine organoid lines from WT and *Sf3b1^K700E/+^*animals were established as previously described (Boj et al., 2015). Briefly, 43-week-old animals were euthanized and their pancreata excised. The organs were dissected to thin pieces and digested in 4 mg/ml collagenase IV for 7 minutes at 37^°^C. Then, pancreatic ducts were manually picked under a light microscope and seeded in drops of growth factor reduced matrigel. In vitro activated (pre-) cancer organoid lines (*Kras^G12D/+^*, *Kras^G12D/+^; Sf3b1^K700E/+^*, *Kras^G12D/+^*; *Trp53^R172H/+^* and *Kras^G12D/+^*; *Trp53^R172H/+^*; *Sf3b1^K700E/+^*) were established from 8-week-old animals using the same protocol, except that recombination was achieved by delivering Cre-GFP by lentiviral transduction, followed by FACS sorting for GFP positive cells. Tumor-derived KPC and KPC-Sf3b1^K700E/+^ organoid lines were established from solid tumors of KPC or KPC-Sf3b1^K700E/+^ mice. Tumor tissue was digested for 2–3 hours in 4 mg/ml collagenase IV at 37^°^C, pelleted and seeded in drops of matrigel. The presence of the K700E mutation was validated with Sanger-sequencing on RNA level for each organoid line. Each organoid line was isolated from an individual mouse. Tumor-derived KPC and KPC-Sf3b1^K700E/+^ organoid lines were additionally plated in regular cell culture dishes and grown as monolayer cell culture. Organoids were cultured in organoid medium (OM) composed of AdDMEM/F12 (Gibco) supplemented with GlutaMAX (Gibco), HEPES (Gibco), Penicillin-Streptomycin (Invitrogen), B27 (Gibco), 1.25 mM N-Acetyl-L-cysteine (Sigma), 10 nM Gastrin I (Sigma) and the growth factors: 100 ng/ml FGF10 (Peprotech), 50 ng/ml EGF (Peprotech), 100 ng/ml Noggin, 100 ng/ml RSPO-1 (Peprotech), and 10 mM Nicotinamide (Sigma). For the first week after duct isolation the culture medium was supplemented with 100 μg/ml Primocin (InvivoGen).

### Immunohistochemistry

Murine tissue specimens were dissected and fixed in 10% neutral buffered formalin for 48 - 72 hours. Thereafter, formalin was replaced with 70% ethanol before paraffin-embedding and sectioning at a thickness of 4 μm. Hematoxylin and eosin stainings were performed according to the manufacturer’s instructions. Masson-Goldner-Trichome staining was performed as previously described (Goldner, 1938). Quantification of the collagenous tissue area was performed with QuPath-0.2.3. The pixel classifier was trained to separate tissue areas stained red from tissue areas stained green and from areas without tissue. Anti-Ki-67 (Antigen Clone TEC-3) antibody (Dako), anti-Cleaved Caspase 3 (Asp175) antibody (Cell Signaling Technology), anti-E-Cadherin (Clone 36) antibody (BD Biosciences), anti-FN-1 antibody (Chemicon) and anti-TNC (Clone 578) antibody was used according to the manufacturer’s recommendations. TNC staining was quantified by calculating the average of Raw Int Density of 3 randomly chosen fields per specimen using ImageJ. Luminal necrotic cells were defined as shed cells residing within the lumen of PanIN-lesions.

### Organoid growth assay

Growth of organoids was assessed with CellTiter-Glo 3D (Promega). For absolute quantification of ATP levels, standard curves with defined concentrations of ATP were used for every measurement according to the manufacturer’s instructions. As approximation of proliferation rate, the ratio of ATP concentrations at the indicated time points was calculated.

### Crystal violet assay

To measure proliferation of cell lines, 5000 cells were seeded per 96 wells. At the indicated time points, cells were stained with 0.5% (w/v) crystal violet (Sigma-Aldrich) dissolved in an aqueous solution with 20% Methanol (v/v). After washing, plates were allowed to air-dry and the crystal violet was dissolved in 10% acetic acid. Optical density was measured at 595 nM in an Infinite 200 plate reader (Tecan).

### Organoid viability assay

For short-term treatment, organoids were seeded as fragments in 10 μl of Matrigel and allowed to form spheres for 24 hours in regular organoid medium. Organoids were thereafter exposed to the indicated concentration of TGF-β1 (Thermo Fisher). To assess viability of the organoids, intact organoids were counted and compared to untreated organoids after 48h of TGF-β1-exposure. This method of quantification was validated by correlating counts of intact organoids with ATP levels as described above (data not shown). For long-term treatment, organoids were seeded as fragments in 40 μl of Matrigel. After allowing to form spheres for 24 hours, the indicated concentration of TGF-β1 was added. After 4 days of TGF-β1-treatment, matrigel-drops were imaged and the number of intact organoids was counted. Then, organoids were reseeded as fragments in normal organoid medium and TGF-β1 was added after 24 hours. Every 4^th^ passage, organoids were split in a 1:1 ratio in the 1ng/ml TGF-β1 condition. Commercial cell lines were seeded and TGF-β1 was added at the indicated concentrations after 12h after plating.

### Organoid invasion assay

Organoids were seeded at equal density in 40 μl of matrigel in 24-well plates. 24 hours after seeding, organoid growth medium was supplemented with 10 ng/ml TGF-β1 (Thermo Fisher). 96 hours after seeding, matrigel domes were detached by rinsing and the migrated organoids (i.e. cells attached to the cell culture dish) were stained by crystal violet. The fraction of attached organoids was calculated by dividing the number of attached organoids by the number of attached organoids plus the number of non-attached organoids (i.e. organoids residing in the matrigel dome).

### Cleaved-caspase 3/7 assay

Organoids were seeded 10 μl of Matrigel and allowed to form spheres for 24 hours in regular organoid medium. Organoids were thereafter exposed to 10 ng/ml TGF-β1 (Thermo Fisher) overnight. Cleavage of Caspase 3 and 7 was quantified by using Caspase-Glo 3/7 Assay System (Promega) according to the manufacturer’s instructions.

### Chemical inhibitors

The following chemical inhibitors targeting different effectors of the TGF-β-pathway were used: TGFbR-inhibitor A83-01 [50 nM] (Tocris Bioscience), p38-inhibitors SB202190 [10 μM] (Sigma-Aldrich) and SB203580 [10 μM] (Selleckchem), JNK-inhibitor SP600125 [25 μM] (Sigma-Aldrich), SMAD3-inhibitor SIS3 [10 μM] (Sigma-Aldrich) and MAP3K7-inhbitor Takinib [5 μM] (Sigma-Aldrich). The inhibitors were added to the organoid medium directly after seeding.

### shRNA-mediated *Map3k7* knockdown

shRNA targeting murine *Map3k7* was purchased from Sigma-Aldrich (TRCN0000022563). A pLKO.1-puro Non-Target shRNA was used as control. Lentivirus was produced by PEI-based transfection of HEK293T cells. Briefly, HEK293T cells were seeded at 70% confluency in 6-well plates, and the following plasmids were transfected: PAX2 plasmid (1100 ng), VSV-G plasmid (400 ng), cargo plasmid (1500 ng). Medium was changed 12 hours after transfection and the virus-containing supernatant collected after 36 hours. Organoids were dissociated into single cells by Tryp-LE treatment for 5 minutes at 37^°^C and consecutive mechanical disruption. After centrifugation, 10% Lentivirus-containing supernatant in organoid medium (v/v) was added to the cell suspension. After a 4–6h incubation at 37^°^C, cells were seeded in matrigel as described above. Organoids were selected in 2ng/ml Puromycin after the first passage for at least 5 days.

### Overexpression of *Map3k7*

Murine *Map3k7* (full-length isoform) was amplified from cDNA of murine WT duct organoids and cloned into a Lenti-backbone. Production of lentivirus and transduction of organoids was performed as described above. Organoids stably overexpressing GFP (addgene #17488) were used as experimental control.

### Overexpression of SF3B1-K700E

Codon-optimized human SF3B1-WT and SF3B1-K700E was derived from the plasmids pCDNA3.1-FLAG-SF3B1-WT (addgene #82576) and pCDNA3.1-FLAG-hSF3B1-K700E (addgene #82577) and cloned into a Lenti-backbone. The lentiviruses were produced as described above and used to transduce various cell lines. Puromycin-selection was used to select for transduced cells for at least two passages.

### RNA sequencing

#### Cell sorting and RNA extraction

Murine tumors were excised and digested for 2–3 hours in collagenase (4mg/ml) at 37^°^C. After addition of fetal calf serum (FCS) to stop the digestion, cells were strained trough a 100 μm and a 70 μm cell strainer. Then, cells were washed twice in PBS + 2% FCS + 2 mM EDTA and incubated with mouse FcBlock (BD Biosciences), Epcam-APC (CD326 Monoclonal Antibody (G8.8), APC, eBioscience™, Thermo Fisher) and CD45-BV785 (Clone 30-F11, Biologened) antibodies for 30 minutes at 4^°^C. After washing, Epcam-positive-CD45-negative cells were sorted into lysis buffer with a BD FACSAriaIII Cell Sorter (BD Biosciences). Finally, RNA was extracted using NucleoSpin RNA XS kit (Macherey Nagel) according to the manufacturer’s instructions.

### Library preparation

The quantity and quality of the isolated RNA was determined with a Qubit® (1.0) Fluorometer (Life Technologies, California, USA) and a Tapestion (Agilent, Waldbronn, Germany). The SMARTer Stranded Total RNA-Seq Kit - Pico Input Mammalian (Clontech Laboratories, Inc., A Takara Bio Company,California, USA) was used in the succeeding steps. Briefly, total RNA samples (0.25–10 ng) were reverse-transcribed using random priming into double-stranded cDNA in the presence of a template switch oligo (TSO). When the reverse transcriptase reaches the 5’ end of the RNA fragment, the enzyme’s terminal transferase activity adds non-templated nucleotides to the 3’ end of the cDNA. The TSO pairs with the added non-templated nucleotide, enabling the reverse transcriptase to continue replicating to the end of the oligonucleotide. This results in a cDNA fragment that contains sequences derived from the random priming oligo and the TSO. PCR amplification using primers binding to these sequences can now be performed. The PCR adds full-length Illumina adapters, including the index for multiplexing. Ribosomal cDNA is cleaved by ZapR in the presence of the mammalian-specific R-Probes. Remaining fragments are enriched with a second round of PCR amplification using primers designed to match Illumina adapters.The quality and quantity of the enriched libraries were validated using Qubit® (1.0) Fluorometer and the Tapestation (Agilent, Waldbronn, Germany). The product is a smear with an average fragment size of approximately 360 bp. The libraries were normalized to 10nM in Tris-Cl 10 mM, pH8.5 with 0.1% Tween 20.

### Cluster Generation and Sequencing

The TruSeq SR Cluster Kit HS4000 or TruSeq PE Cluster Kit HS4000 (Illumina, Inc, California, USA) was used for cluster generation using 8 pM of pooled normalized libraries on the cBOT. Sequencing was performed on the Illumina HiSeq 4000 paired end at 2 X126 bp or single end 126 bp using the TruSeq SBS Kit v4-HS (Illumina, Inc, California, USA).

### RNAseq data analysis

Adapters have been trimmed with trimmomatic (v0.35). Pairs for which both reads passed the trimming have been mapped to the murine genome using STAR (v2.7.0a) and indexed BAM files obtained with samtools (v1.9). Reads were counted with featureCounts from subread package (v1.5.0). The read counts have been processed in a statistical analysis using edgeR (v3.24.3), obtaining a list of genes ranked for differential expression by p-value and Benjamini-Hochberg adjusted p-value as the estimate of the false discovery rate. All data is summarized in Table 1.

### Gene set enrichment analysis

Gene set enrichment analysis (v.4.1.0, Broad Institute, MIT) was used to determine enriched gene sets in KPC or KPC-Sf3b1^K700E/+^ tumor cells. Standard parameters of the software were used to perform the analysis. Molecular Signatures Database v7.4, Hallmark Gene Sets (H) was used to query enriched gene sets. The input gene expression matrix contained read-count information (count per million) of 21,633 genes.

### Alternative splicing analysis

We ran a 2-pass alignment of the fastq files using STAR v2.7 (Dobin et al., 2013) using the GRCm38.p6 genome as reference. The gene annotation used was GENCODE v.m25. For gene expression quantification we used a custom script, available at github: https://github.com/ratschlab/tools-omicstools/tree/master/gromics/counting; commit hash d074114f1d0a9f518c9cd039f68de0cdf8d583ff.

SplAdder v.2.2 (Kahles et al., 2016) was run to build splicing graphs and determine splice events. Differential splicing events were determined by calculating a log(psi+x) transformation of the percent spliced in (calculated as ratio of reads supporting the splice event over the number of reads supporting the alternate event). Splice events that did not show any variability over the samples were removed and missing values were mean imputed. After standardization a two-sided t-test was used to calculate p-values of splice events differences between KPC and KPC-Sf3b1^K700E^ mice. All data is summarized in Table 2.

### Motif analysis

Consensus 3’ ss motif in proximity of the canonical and the cryptic 3’ ss in sorted KPC-Sf3b1^K700E/+^ tumor cells was assessed by query 30-40 bases spanning the respective 3’ss of the 7 main splice events for a motif using weblogo-sequence creator (https://weblogo.berkeley.edu/logo.cgi).

### RT-PCR and quantitative RT-PCR (qPCR)

RNA-extraction was performed with QIAGEN RNeasy Mini Kit, and cDNA was generated with GoScript Reverse Transcriptase kit (Promega) according to the manufacturers’ instructions. RT-PCR was performed with GoTaq G2 Green Master Mix (Promega) and gene specific primers. Amplicons were fractionated on 2% TBE gel (Life Technologies) supplemented with 0.01% GelRed (Biotium). For qPCR, 2 μL of 1:10-diluted cDNA was added to 8 μl of 5x HOT FIREPol Evagreen qPCR Supermix (SolisBiodyne). RT-qPCR was performed with a LightCycler480 II (Roche). Relative gene expression was determined with the comparative CT method. Genes with a median CT value of more than 33 cycles and a difference of less than 3.3 cycles to the template control (H_2_O) were defined as not detectable. Sequences of all primers used in this study are listed in Supplementary Table 1.

### NGS-based isoform quantification of *Map3k7*

Primers generating an amplicon including the exon 4 and exon 5 junction of Map3k7 cDNA were used. Briefly, a gene-specific amplicon was generated in a 20 μL reaction for 35 cycles with GoTaq G2 Green Master Mix (Promega). The PCR product was purified using the NucleoSpin Gel and PCR Clean-up kit (Macherey-Nagel). Thereafter, the isolated product was amplified for 8 cycles using primers with sequencing adapters. After column-based isolation of the amplicon and quantification of DNA-yield using a Qubit 3.0 fluorometer and the dsDNA HS assay kit 392 (Thermo Fisher), paired-end sequencing was performed on an Illumina Miseq. The sequencing data was subsequently analyzed with CRISPResso2 (Clement et al., 2019).

### Western blotting

Cells were lysed in RIPA buffer, supplemented with Protease Inhibitor (Cell Signaling Technologies) and PhosStop (Sigma-Aldrich) and centrifuged for 10 min at 21’000 g. The protein concentration of the supernatant was determined using Pierce BCA assay (ThermoFisher) and a standard curve of albumin. Then, samples were heated for 5’ at 95^°^C in Lämmli buffer and protein lysates were resolved on polyacrylamide Mini-PROTEAN TGX gels (BioRad) and transferred onto nitrocellulose membrane by wet-transfer. The following antibodies were used for immunoblotting: Recombinant anti-GADPH (EPR16891, Abcam) and rabbit monoclonal anti-MAP3K7 (anti-TAK1, D94D7, Cell Signaling Technology) IRDye-conjugated secondary antibodies (donkey anti-goat: LI-COR cat. no. 926-32214; anti-rabbit: LI-COR cat. no. 926-68073) were used for signal detection by an Odyssey Imager (LI-COR) imaging system.

### Data availability

The RNA sequencing raw data were deposited in the NCBI Gene Expression Omnibus (GEO) under accession number GSE203339. Splice analysis of human cancers was performed on a previously published dataset, accessible at https://gdc.cancer.gov/about-data/publications/PanCanAtlas-Splicing-2018 (Kahles et al., 2018). Material created in this study (i.e. primary cell lines, pasmids) are provided upon request and shall be directed at the corresponding author of this study (Prof. G. Schwank). Source data of the blots shown in Fig. 4E,F, Fig. 5D and Suppl. Fig. 4D,E,G can be found in source file zip in this manuscript.

## SUPPLEMENTAL INFORMATION

**Supplementary Figure 1.**
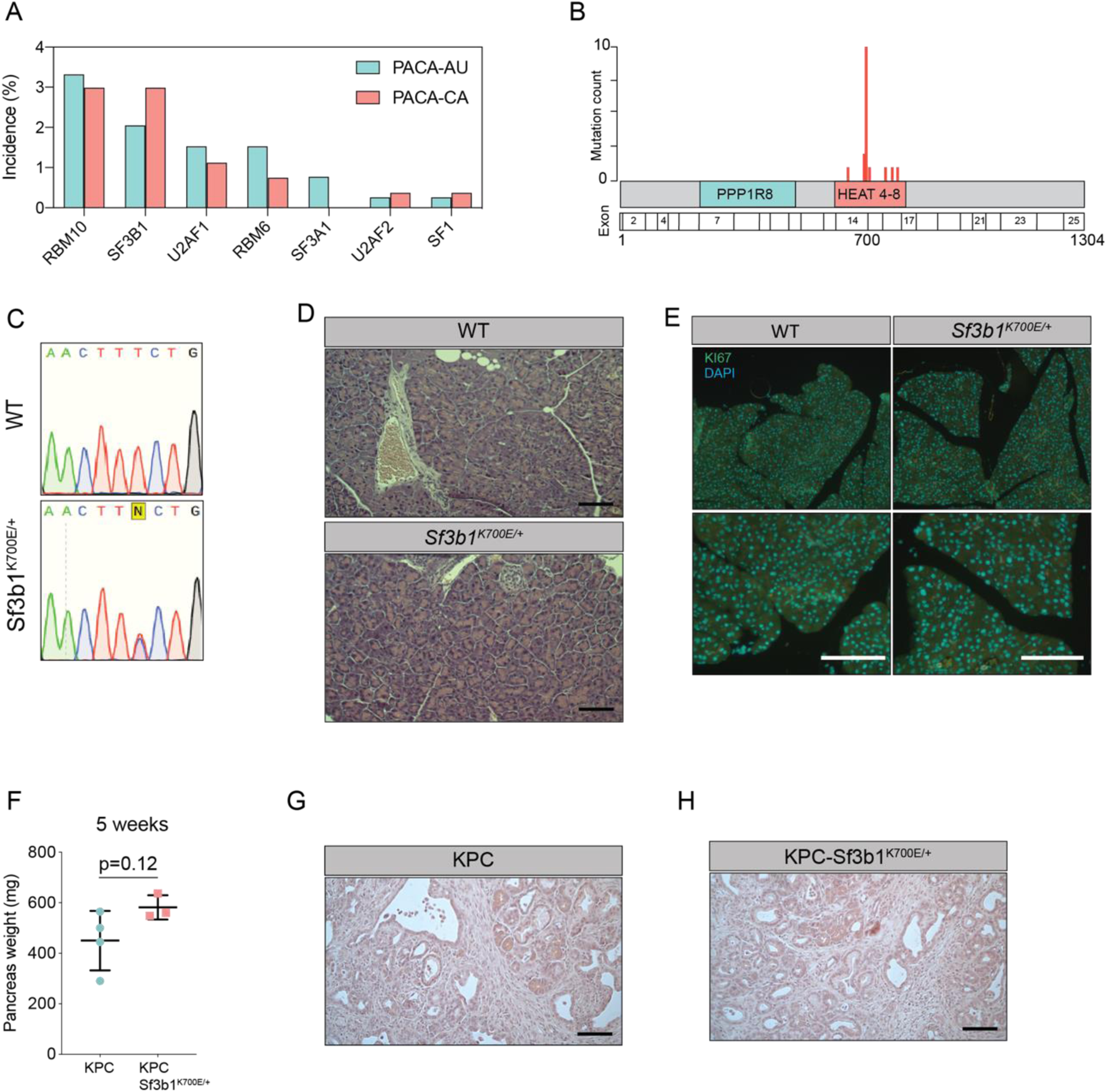
**(A)** Incidence of mutations in splicing factors in PDAC patients derived from the ICGC database (PACA-AU, n=391 and PACA-CA, n=268). **(B)** Incidence of *SF3B1* missense mutations in PDAC patients derived from PACA-AU and PACA-CA. **(C)** Representative Sanger-sequencing results of the *Sf3b1^K700E/+^* mutation (T>C) of cDNA isolated from pancreata at 43 weeks of age of *Ptf1a-Cre* (WT) or *Ptf1a-Cre; Sf3b1^K700E/+^* (*Sf3b1^K700E/+^*) mice. **(D-E)** WT and Sf3b1^K700E/+^ pancreata at 43 weeks of age stained with H&E (D) and Ki67 (E) Scale bar is 50 μm. **(F)** Pancreatic weight of KPC and KPC-Sf3b1^K700E/+^ mice, at 5 weeks of age. Two-tailed unpaired t-test was used to compute the indicated p-value. **(G-H)** Representative micrograph images of KPC (G) and KPC-Sf3b1^K700E/+^ (H) pancreata at 9 weeks of age stained with H&E, scale bar is 100 μM.

**Supplementary Figure 2.**
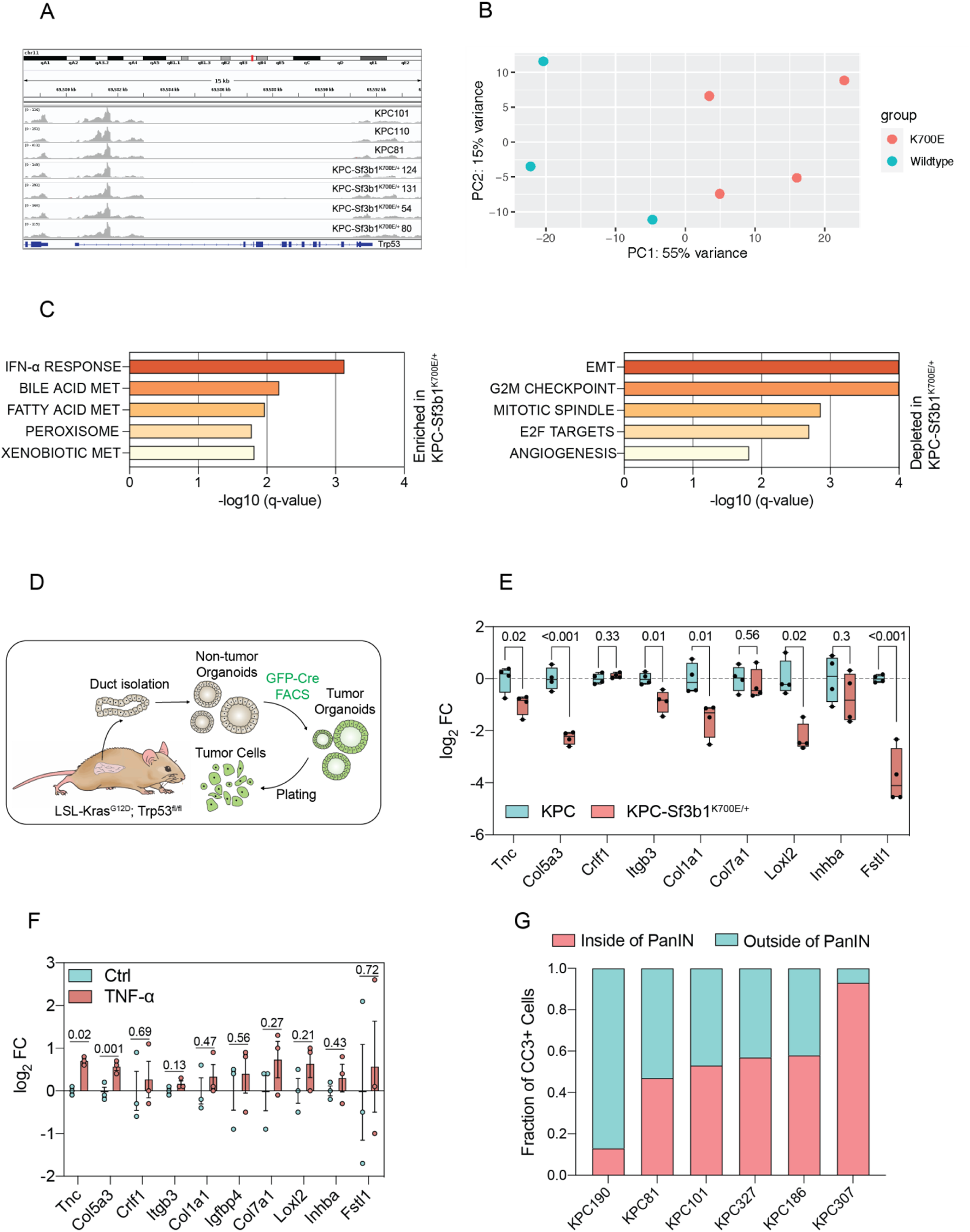
**(A)** Integrated genome viewer (IGV) displaying RNA-seq reads of *Trp53* of sorted KPC and KPC-Sf3b1^K700E/+^ cells. **(B)** Principal component analysis showing the variance in two dimensions in relation to the genotypes of the sorted tumor cells (Wildtype = KPC, K700E = KPC-Sf3b1^K700E/+^). **(C)** Results of GSEA analysis, displaying most enriched (top) and depleted (bottom) GSEA-Hallmark pathways in KPC-Sf3b1^K700E/+^ animals. **(D)** Schematical overview of generation of in-vitro activated tumor cells: Pancreatic ducts of LSL-*Kras^G12D/+^; Trp53^fl/f^* or LSL-*Kras^G12D/+^; Trp53^fl/fl^; Sf3b1^flK700E/+^* were isolated and expanded as organoids. Cells were transduced with GFP-Cre and selected by FACS. After expansion of sorted cells in a 3D culture, organoids were plated in cell culture dishes and grown as monolayer culture. **(E)** RT-qPCR analysis of EMT genes displayed in (G) of KPC (n=3) and KPC-Sf3b1^K700E/+^ (n=3) in-vitro activated cancer cell lines treated with TGF-β1 (10 ng/ml) for 24 hours. The experiment was performed independently 4 times for every cell line, the average of all replicates is shown. *Col3a1, Sfrp1, Igfbp4, Col1a2, Mmp2* and *Lama1* were not detected and therefore excluded from analysis (see methods for details). Two-tailed unpaired t-test was used to compute the indicated p-values. **(F)** RT-qPCR analysis of EMT genes displayed in Fig 2B in 3 different KPC cell lines treated with TNF-α (100 ng/ml) for 24 hours. The experiment was performed independently 3 times for every cell line. *Col3a1, Sfrp1, Igfbp4, Col1a2, Mmp2* and *Lama1* were not detected and therefore excluded from analysis (see methods for details). **(G)** Quantification of CC3 posivtive (CC3+) cells residing within PanIN lesions in 6 different KPC tumor specimen.

**Supplementary Figure 3.**
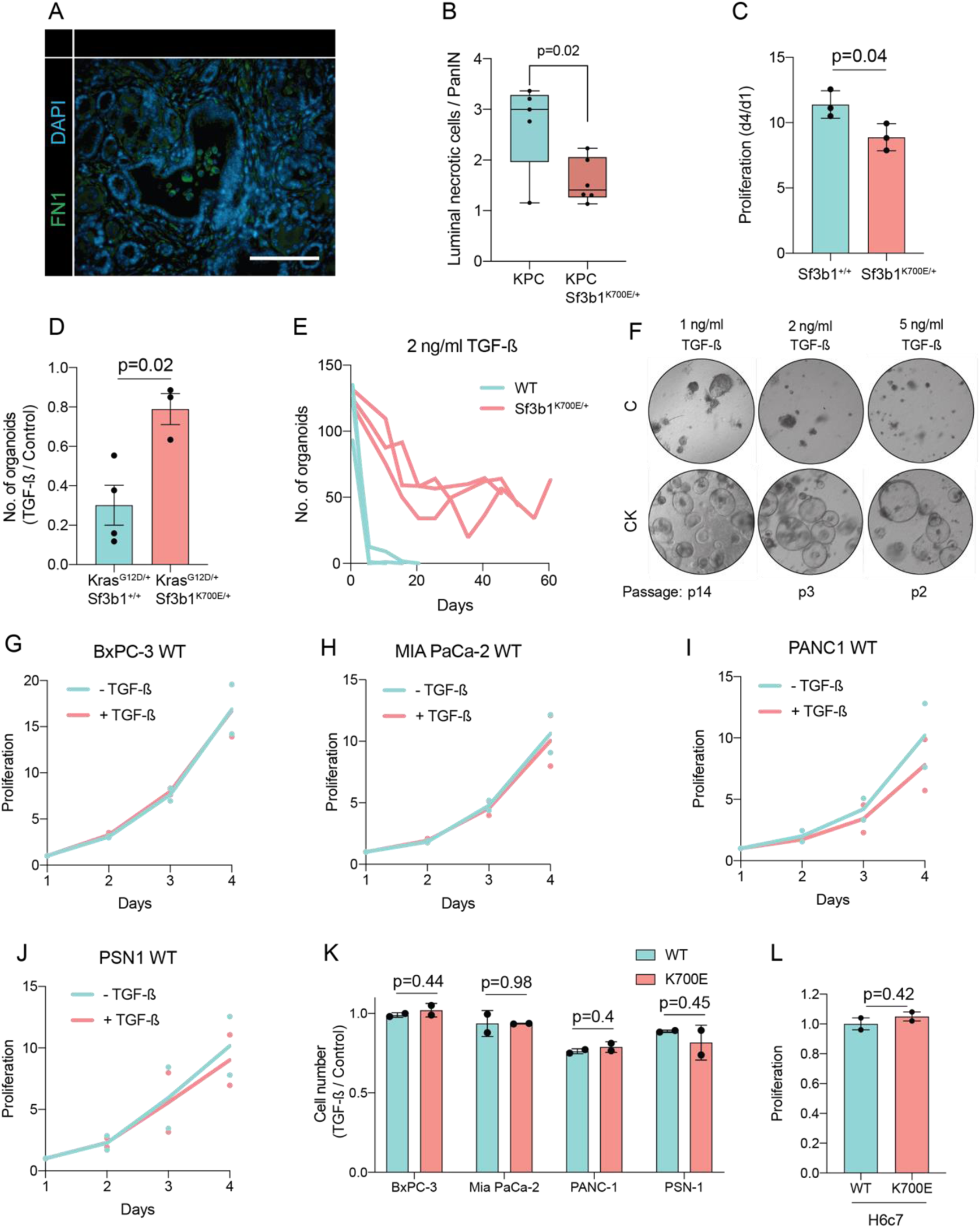
**(A)** Representative microscopy images of FN1 (green) in murine PDAC. Scale bar is 50 μm. **(B)** Blinded quantification of luminal necrosis in KPC (n=5) and KPC-Sf3b1^K700E/+^ (n=6) tumor samples. The average number of necrotic cells per PanIN lesion is plotted, two-tailed unpaired t-test was used to compute the indicated p-value. **(C)** Proliferation of pancreatic ductal organoids derived from WT and *Sf3b1^K700E/+^*mice without TGF-β1 supplementation. Two-tailed unpaired t-test was used to compute the indicated p-value. **(D)** Organoid count of organoids of the indicated genotypes exposed to 10 ng/ml TGF-β1 for 48h. Two-tailed unpaired t-test was used to compute the indicated p-value. **(E)** Organoid count of organoids cultured in medium containing 2 ng/ml TGF-β1 for the indicated period of time. One organoid line for each genotype was used, the experiment was independently performed twice. **(F)** Representative microscopy images of WT and *Sf3b1^K700E/+^* organoids exposed to 1, 2 or 5 ng/ml TGF-β1 at the indicated number of passages. **(G-J)** Proliferation of the indicated PDAC cell lines overexpressing SF3B1 exposed to 10 ng/ml TGF-β1 compared to normal growth medium. The experiment was independently performed twice. **(K)** Viability of indicated cell lines overexpressing wildtype or mutated SF3B1 after 72 hours of exposure to 10 ng/ml TGF-β1. The experiment was independently performed twice, two-tailed unpaired t-test was used to compute indicated p-values **(L)** Normalized growth of the pancreatic duct cell line H6c7 overexpressing SF3B1-WT or SF3B1-K700E after 4 days of culture in normal growth medium. The experiment was independently performed twice, two-tailed unpaired t-test was used to compute indicated p-value.

**Supplementary Figure 4.**
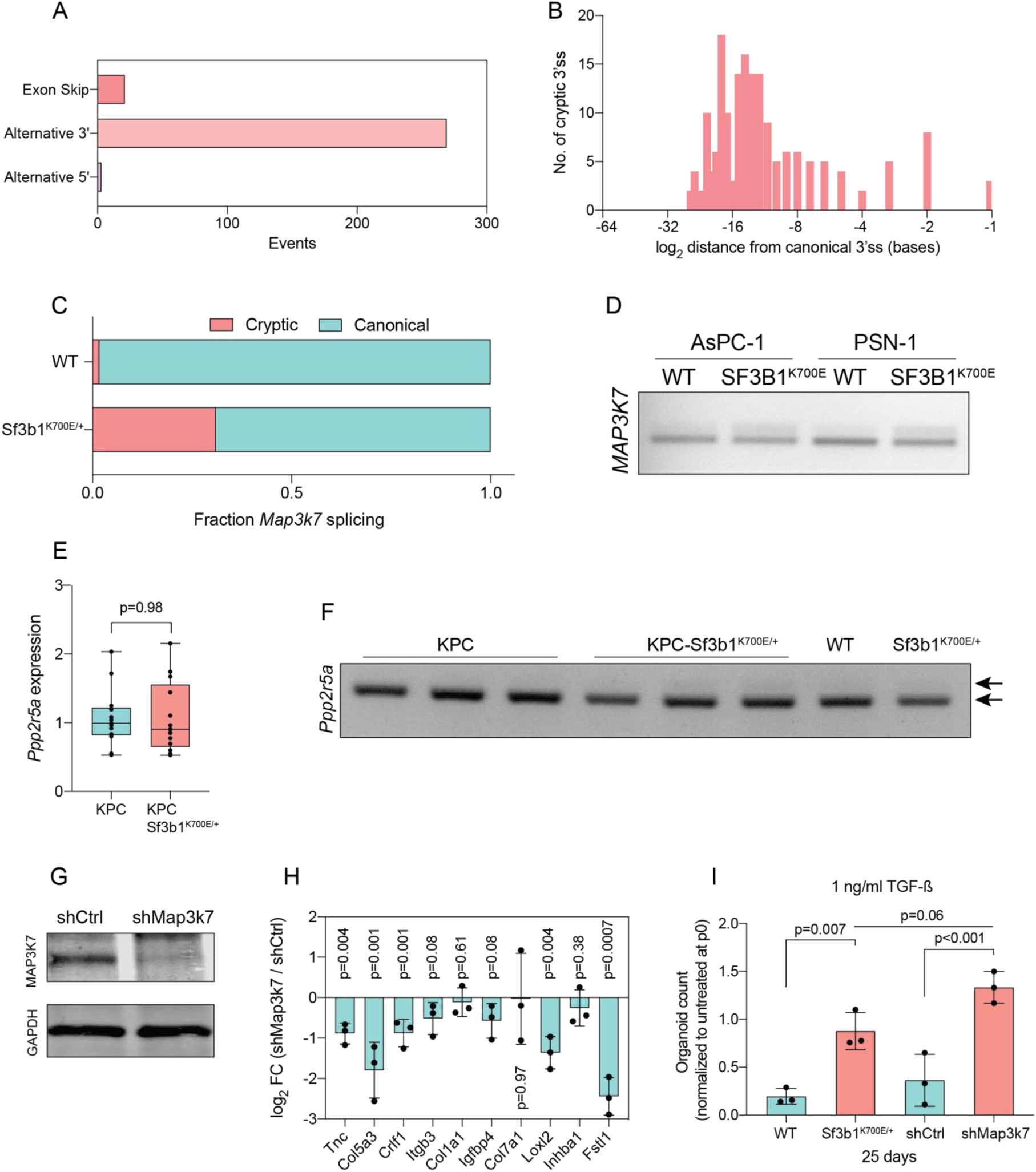
**(A)** Pan-cancer analysis of alternative splice-events identified in solid tumors carrying the *SF3B1^K700E^* mutation (PSI>0.05, FDR<1^-10^). **(B)** Histogram displaying the distance of cryptic 3’ss from the adjacent canonical 3’ss on a logarithmic scale in solid tumors carrying the *SF3B1^K700E^*mutation. **(C)** NGS-results of *Map3k7* cDNA isolated from WT and *Sf3b1^K700E/+^* pancreas organoids (n=1). **(D)** RT-PCR amplicon of *MAP3K7* cDNA isolated from four human PDAC cell lines overexpressing wildtype SF3B1 (OE WT) or K700E-mutated SF3B1 (OE K700E). The amplicon includes the 3’ splice site of exon 4 and 5, the upper band of the gel image represents the non-canonical transcripts. **(E)** RT-qPCR of *Ppp2r5a* expression in KPC (n=13) and KPC-Sf3b1^K700E/+^ (n=12) cancer-derived organoid lines. Two-tailed unpaired t-test was used to compute the indicated p-value. **(F)** RT-PCR amplicon of *Ppp2r5a* cDNA isolated from sorted KPC (n=3) and KPC-Sf3b1^K700E/+^ (n=4), as well as WT and *Sf3b1^K700E/+^* pancreas organoids. The amplicon includes the 3’ splice site of exon 4 and 5, the upper arrowhead represents the predicted size of non-canonical transcripts. **(G)** Western blot gel-image of MAP3K7 and GAPDH in a KPC cell line transduced with a control shRNA (shCtrl) or a shRNA targeting Map3k7 (shMap3k7). **(H)** RT-qPCR analysis of EMT genes displayed in Fig. 2G in a KPC cell line transduced with a control shRNA (shCtrl) or a shRNA targeting Map3k7 (shMap3k7) treated with TGF-β1 (10 ng/ml) for 24 hours. The experiment was performed independently 3 times. **(I)** Organoid count normalized to untreated organoids of respective genotypes at passage 0. Duct organoids of indicated genotypes / treated with shRNA were exposed to 1 ng/ml TGF-β1 for 25 days. Data represents one organoid line per condition, the experiment was independently performed three times. One-way ANOVA was used to compute the indicated p-values.

